# Testing for age- and sex- specific mitonuclear epistasis in *Drosophila*

**DOI:** 10.1101/2024.10.10.617520

**Authors:** Martin D. Garlovsky, Ralph Dobler, Ruijian Guo, Susanne Voigt, Damian K. Dowling, Klaus Reinhardt

## Abstract

The need for efficient ATP production is predicted to result in the evolution of cooperation between the mitochondrial and nuclear encoded components of the electron transport system. Novel (i.e., mismatched) mitonuclear genotype combinations are therefore predicted to result in negative fitness consequences, which may become more prevalent with ageing. Such negative fitness effects are expected to be prominent in males, since maternal inheritance of mitochondria is predicted to lead to accumulation of male-harming mutations (the mother’s curse hypothesis). To test these predictions, we measured female and male fertility traits using a genetically diverse panel of 27 mitonuclear populations of *Drosophila melanogaster* with matched or experimentally mismatched mitonuclear genomes at different ages. We found no overall effect of mitonuclear mismatch. In females, we found no effect of mitonuclear epistasis. In males, we found limited evidence of mitonuclear epistasis affecting fitness in old age, however, not in the direction predicted. Experimentally mismatched males sired more offspring in one comparison. Sex-specific advantages of mismatched males might arise if novel nuclear alleles compensate for deleterious mitochondrial alleles that have accumulated. If such compensatory effects of novel mitonuclear combinations increasing fitness occur in nature, they could represent a possible counterforce to the mother’s curse.

## INTRODUCTION

Mitochondria provide energy via ATP production in most eukaryotic cells. The electron transport system that generates the proton gradient across the inner mitochondrial membrane driving oxidative phosphorylation is encoded across two genomes; the mitochondrial genome and the nuclear genome (Ballard and Rand 2005; St John et al. 2005; Piomboni et al. 2012; Burton et al. 2013; Stojković and Đorđević 2017). Efficient energy production therefore selects for cooperation between mitochondrial and nuclear encoded components of the electron transport system (Rand et al. 2004; Dowling et al. 2008; Lane 2011; Dowling 2014; Hill 2019; Hill et al. 2019).

Studies that have experimentally created novel pairings of mitochondrial and nuclear genotype (hereafter mitochondrial replacement studies) have frequently revealed mitonuclear epistasis, where the effects of a given mitochondrial genotype vary depending on the nuclear background with which it resides and *vice versa* (Immonen et al. 2016a; Đorđević et al. 2017; Hill 2020; Camus et al. 2023). Replacing lineage-specific, putatively coevolved (i.e., “matched”) mitonuclear genotypes with novel (i.e., “mismatched”) mitonuclear combinations has revealed considerable fitness effects, however the direction may often be unpredictable (reviewed in (Dobler et al. 2018). Many studies have shown mismatched mitonuclear genotypes result in reduced fitness, e.g., reduced lifespan, neuronal and metabolic activity, sperm competitive ability, and increased incidence of disease (Ellison and Burton 2008; Smith et al. 2010; Aw et al. 2011; Reinhardt et al. 2013; Yee et al. 2013, 2015; Dobler et al. 2014, 2018; Wolff et al. 2016; Zhang et al. 2017; Vaught and Dowling 2018). However, novel mitonuclear genotypes have also been suggested to offer the potential to be advantageous (reviewed in (Eyre-Walker 2017).

Negative epistatic fitness effects exposed by mitonuclear mismatch are thought to arise because mitochondrial variants that work efficiently in the nuclear background in which they have (co-)evolved may be deleterious in a novel genetic background (Dobler et al. 2018; Hill et al. 2019; Hill 2020). For instance, because of the maternal inheritance of mitochondria (Birky 1978), selection will be ineffective at purging male-deleterious mitochondrial mutations if they have a neutral or positive effect in females. Female beneficial/male deleterious alleles can therefore accumulate within populations, resulting in a sex-biased mutation load, sometimes termed the ‘mother’s curse’ effect (Frank and Hurst 1996; Gemmell et al. 2004; Unckless and Herren 2009; Smith et al. 2010; Innocenti et al. 2011; Dobler et al. 2014; Wolff et al. 2016; Milot et al. 2017; Nagarajan-Radha et al. 2020; Camus et al. 2023). These male-deleterious mitochondrial mutations are predicted to select for counteradaptations (rescue mutations) in the nuclear genome to offset the harmful mitochondrial effects. Mitochondrial replacement experiments can expose such harmful phenotypic effects because male-deleterious mitochondrial mutations are placed into a novel nuclear genetic background which lacks the requisite rescue mutations (Hill 2020). Consequently, traits expressed only in males, such as sperm traits, are predicted to be particularly sensitive to mitonuclear interactions (Frank and Hurst 1996) as natural selection has little power to act on mitochondrial variation affecting the expression of sperm traits via the female line (Friesen et al. 2020, Vaught and Dowling 2018). Despite the predicted sex-specific effects of mitonuclear interactions, relatively few studies have investigated male-specific traits (but see (Innocenti et al. 2011; Yee et al. 2013; Immonen et al. 2016b,a; Camus and Dowling 2018; Vaught and Dowling 2018; Keaney et al. 2020).

Reproductive senescence, the decline in reproductive output with increasing age (e.g., (Novoseltsev et al. 2003), has been studied extensively in female animals, with historically less focus on traits that may contribute towards male reproductive senescence (Fricke and Koppik 2019; Fricke et al. 2023). Many mitochondrial effects have age-specific phenotypic effects or disease penetrance (e.g., (Bratic and Larsson 2013; Immonen et al. 2016a; Martikainen et al. 2017; Đorđević et al. 2017; Dunham-Snary et al. 2018), suggesting the cooperation between the mitochondrial and nuclear components of the electron transport system may be less efficient in old age. If, as expected, mitonuclear cooperation is less efficient in old age, then the effects of mitonuclear mismatch will be evident as accelerated reproductive senescence in mismatched, compared to matched mitonuclear combinations. Therefore, mitonuclear interactions affecting sperm function might be more (or only) apparent in old age, where repair machinery and oxygen radical scavengers are less efficient at ameliorating damage (Immonen et al. 2016a; Đorđević et al. 2017).

Here we investigated the effects of mitonuclear mismatch on female and male reproductive senescence using a panel of *Drosophila melanogaster* populations harbouring either matched or experimentally mismatched mitonuclear genotype combinations. The same mitonuclear populations have previously been used to test the effect of mitonuclear epistasis on fecundity, development, and lifespan, separated for immediate and parental effects, or the effect of diet or the epigenomic regulation of the nuclear genome (Grunau et al. 2018; Vaught et al. 2020; Dobson et al. 2023). The three source populations used to establish the mitonuclear replacement panel have diverged genetically, with fixed single nucleotide polymorphisms (SNPs) between populations in both the mitochondrial (Vaught et al. 2020) and nuclear genomes (Dobson et al. 2023). There are 44 segregating SNPs in the mitochondrial genome that differ both within and between mitonuclear populations and significant genetic divergence between populations in the nuclear genome as well as segregating nuclear genomic variation within populations (Dobson et al. 2023). This allowed us to identify mitonuclear variation affecting fitness traits, and further to identify specific mitochondrial haplotypes (“mito-types”) contributing to variation in fitness (Fig. 1).

**Figure 1.**
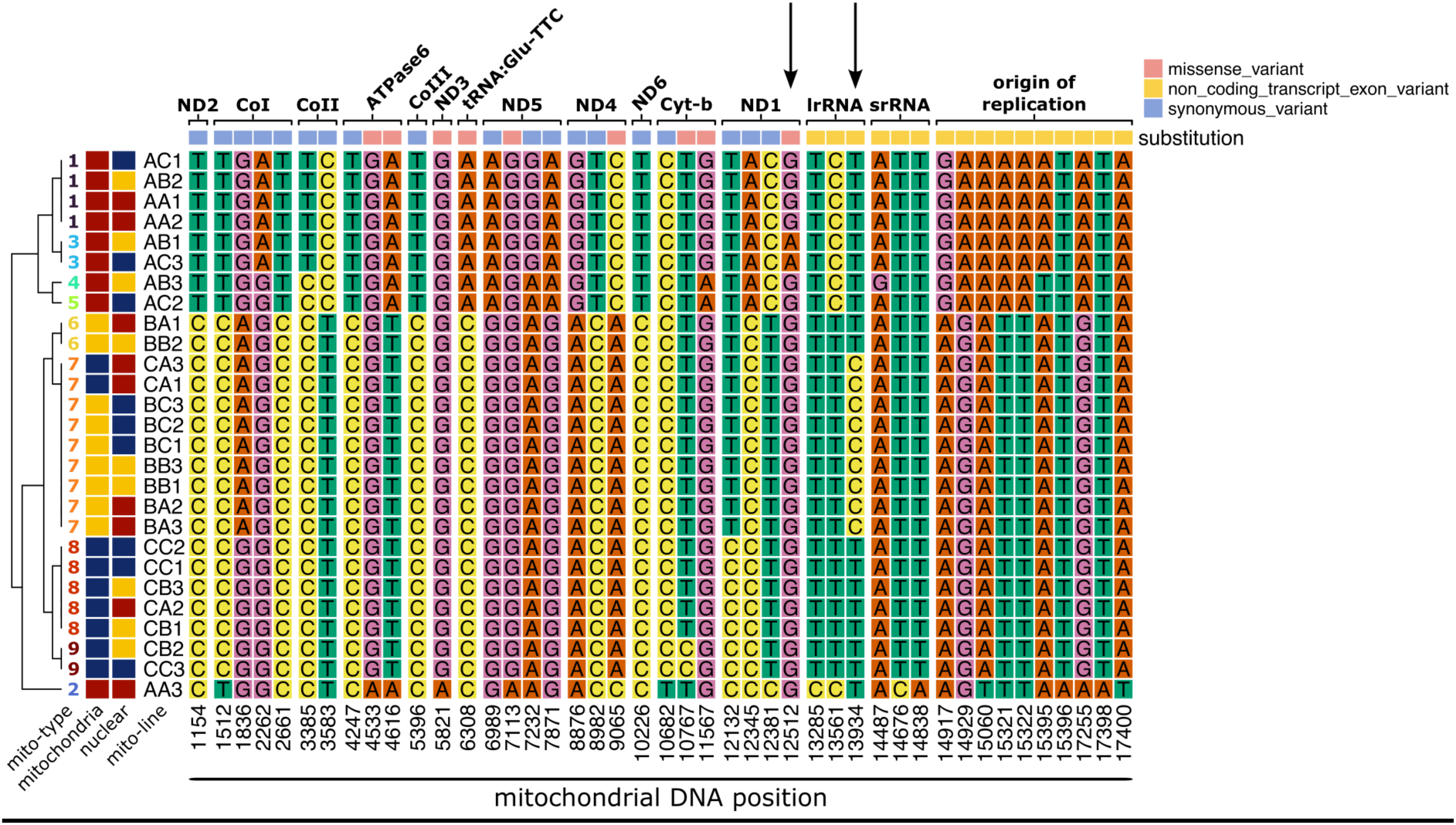
Segregation of major allele frequency differences in the mitochondrial genome between the 27 mitonuclear populations based on significantly differentiated SNPs (Vaught et al. 2020; Dobson et al. 2023). The two-letter notation for each population on the left-hand side denotes the geographic origin of the mitochondrial genotype followed by the nuclear genotype (A: Australia, B: Benin, C: Canada) followed by the replicate population (1-3). The populations cluster into nine hierarchical groups, called 9 ‘mito-types’ denoted in the dendrogram on the far left. The genes containing SNPs are indicated at the top with the variant classification and mitochondrial DNA position along the bottom.

Overall, we predicted higher fitness (measured via reproductive success or expression of reproductive traits) for matched compared with mismatched mitonuclear genotypes. We also predicted more negative fitness effects of mitonuclear mismatch in males compared to females due to the mother’s curse. Finally, the male germline is expected to be better protected from the effects of ageing than the soma (Turnell et al. 2021). Therefore, we predicted stronger negative effects of mitonuclear mismatch with ageing on traits relating to somatic tissue products compared with those produced by the germline (Pizzari et al. 2008; Fricke and Koppik 2019). For instance, males can affect partner fecundity via seminal fluid products secreted from the accessory glands in *Drosophila*, i.e., soma (Immonen et al. 2016a; Fricke et al. 2023), whereas sperm metabolic rate, sperm viability, and hatching success (a proxy for fertilisation success), are all more directly related to sperm cell quality, i.e., the germline.

## METHODS

### Fly stocks and maintenance

Detailed description of the establishment and maintenance of the mitonuclear populations can be found in Vaught et al. (2020). In brief, we used *D. melanogaster* from three source populations collected in Coffs Harbour, Australia (“A” lines) (Williams et al. 2012; Dowling et al. 2014), Benin (“B” lines) (formerly Dahomey) (Clancy 2008) and Dundas, Canada (“C” lines) (MacLellan et al. 2009). Source populations were maintained with a large census population size (N > 1500).

To generate the full reciprocal set of nine mitonuclear genotype combinations, we initially crossed 45 unmated females with the desired mitochondrial genotype (i.e., from a single source population) to 45 males with the desired nuclear genotype. We created three replicate mitonuclear population for each mitonuclear combination, resulting in 27 populations. For all subsequent generations we crossed 45 unmated females from each mitonuclear population with 45 males from the source population with the desired nuclear genotype. This crossing scheme replaces 50% of the remaining nuclear genome from the original female source population with the desired nuclear genome in each generation. After 17 generations, more than 99.99% of the source population nuclear genome is replaced, resulting in the desired mitochondrial haplotype in a novel nuclear genetic background. Matched populations were backcrossed with their own nuclear genome. Populations were continuously backcrossed every generation to avoid coadaptation between the new mitochondrial and nuclear genome combinations, and to ensure that the nuclear background retains variability of the founder populations. We denote mitonuclear populations with the first letter indicating the mitochondrial origin and the second letter the nuclear origin (e.g., AA: A mitochondrial, A nuclear; AB: A mitochondrial, B nuclear, etc.) followed by replicate number (1-3).

Mitonuclear populations were maintained on 14-day non-overlapping generations at 25°C on a 12:12 hour light-dark cycle. We reared flies in 25 mm diameter vials on 7 ml corn-yeast-sugar medium (corn 90 g/L, yeast 40 g/L, sugar 100 g/L, agar 12 g/L, Nipagin 20 mL/L, propionic acid 3 mL/L) with live dry yeast added. At generation 85 we treated all lines against *Wolbachia* infection for three generations with tetracycline added to the food (0.2 g/L solved in 40 mL ethanol). After the treatment we confirmed the successful elimination of *Wolbachia* by PCR (Richardson et al. 2012). Populations were continued to be introgressed as described above. Here we used populations 25 to 50 after *Wolbachia* treatment for the various female and male fitness parameters.

### Experimental tester flies

To control for the effect of mating-partner genotype on reproductive outcomes we mated all focal experimental mitonuclear population individuals to isogenic, outbred ‘tester’ flies. We created tester flies by crossing females and males from two isogenic L_HM_ lines (Chippindale et al. 2001) (unmated females from one line, males from the other line). Offspring were collected prior to mating. Tester males were 4 to 6 days old, tester females aged 3 to 5 days.

### Body size

We collected four females from each mitonuclear population mated with males from their own population and placed each female into individual vials with 7 ml standard medium and *ad libitum* dry yeast and allowed egg laying overnight. The next day we removed the female and kept vials under standard conditions for 14 days, after which we collected all emerging offspring from each vial (a family), anaesthetised them with CO_2_ and placed them in a micro-centrifuge tube at −20°C until measurement. After thawing, we removed wings with forceps and mounted them on double-sided adhesive tape on a sheet of paper. We imaged each wing at 30 × magnification using the built-in camera of a Leica EZ4HD stereoscope and Leica software (LAS x 2.0.0.14332). We measured the area of a polygon defined by landmarks (modified from (Gilchrist and Partridge 1999); Fig. S1) on the right wing of four flies per sex per family (N = 864) using ImageJ (Schneider et al. 2012).

### Female reproductive success

We paired 5-day old unmated focal females (n = 15 per mitonuclear population) individually with a tester male in vials (“vial 1”) containing 5 mL of standard food and *ad libitum* dry yeast to allow mating. We removed males after 24 hours and left females in vials for another 60 hours to lay eggs. We transferred females to new food vials every 84 hours (3.5 day) for a total of 21 days, after which females were discarded. We kept oviposition vials at 25°C on a 12:12 hour light-dark cycle and counted the total numbers of emerging offspring in each vial after 11 days, and again after 15 days (Fig. S2). With this protocol we counted the total number of adult progeny each female produced: i) in each vial ii) combined over the first two vials (7 days) – before the onset of reproductive senescence (Novoseltsev et al. 2003), hereafter ‘early-life fecundity’; and iii) over the course of 21 days, hereafter ‘lifetime fecundity’.

### Male reproductive success

We measured traits in males aged either 5 days (hereafter ‘young males’) or 6 weeks (hereafter ‘old males’). For sperm competition assays (P1 and P2), we only measured young males due to logistical constraints.

#### Male progeny and fertilisation success

We measured partner fecundity as the numbers of eggs laid by tester females mated to focal males, and hatching success as the proportions of eggs from which offspring emerged. Focal males were paired individually with 5-day old tester females in food vials and left to mate for 24 hours (n = 15 per mitonuclear population and age class). After 24 hours, males were discarded, and females were transferred individually to new food vials every 24 hours for three days. We counted the numbers of eggs laid across the three successive vials and maintained vials at 25°C for seven days and subsequently counted the numbers of eggs from which larvae hatched. Counts of hatched and unhatched eggs were performed blind to male age and mitonuclear population.

#### Sperm viability and sperm quality

We measured sperm quality defined as the difference in sperm viability before (t0) and after exposure (t30) to an osmotic stressor medium (Eckel et al. 2017). For each focal male (n = 15 per mitonuclear population and age), the paired seminal vesicles were dissected in 10 µL of phosphate buffer saline (PBS) on a glass slide. To measure sperm viability, one vesicle was transferred to a new slide and sperm released into 10 µL PBS by puncturing the seminal vesicle with a fine insect pin and spreading the vesicle contents evenly by gently shaking the slide horizontally. The sample was immediately stained with live/dead kit (Invitrogen), i.e., 0.5 µL SYBR14® and 1 µL propidium iodide. The second vesicle from each male was transferred into 10 µL of stress buffer (PBS:distilled H2O 1:1.5) on a second glass slide and vesicle contents spread as above. Sperm were exposed to the stress buffer in a moisture chamber for 30 minutes before live/dead staining as above (termed t30). After staining, sperm samples were covered with a cover slip and imaged using an epifluorescence microscope (Leica DMi8, Germany) without incubation (Eckel et al. 2017). For each sample, five areas were haphazardly selected, and images taken at 400 × magnification. The numbers of live (green) and dead (red) sperm were counted in ImageJ V. 1.34 (Schneider et al. 2012).

#### Sperm metabolic rate

We measured the metabolic rate of sperm dissected from seminal vesicles using NAD(P)H autofluorescence lifetime imaging (FLIM) (Wetzker and Reinhardt 2019). For details see the online supplementary material. The entire reproductive tract including seminal vesicles of each male was dissected in 20 µL of PBS on a glass slide (n = 5 per mitonuclear population). The reproductive tract was transferred to a second 20 µL drop of PBS and covered with 22 × 22 mm cover slip with clay spacers at the corners to avoid rupturing the seminal vesicles and sealed with nail polish before FLIM analysis.

#### Sperm competition – offence (P1) and defence (P2)

Brown eyed tester females and competitor males were generated by crossing females and males from two isogenic L_HM_ lines homozygous for a recessive brown eye mutation, using different lines for females and males. Tester females, competitor males and focal males were collected within 6 hours of eclosion and kept in single sex groups of 20 until mating. We assessed 45 focal males per mitonuclear population (total = 1215 focal males) split across three blocks with a random subset of 15 males from each population (i.e., 405 males) per block. Each block was separated by two generations.

On the morning of day 1, we placed tester females in a food vial with a focal or competitor (brown eyed) 5-day old male and allowed the pair the opportunity to mate overnight for 24 hours at 25°C on a 12:12 hour light:dark cycle. We removed males the following morning and kept females in the first vial for 24 hours. On the morning of day 3 we added a competitor (for P1) or focal male (for P2) and recorded the start and end of all copulations to confirm females remated and record copulation duration. If females did not remate within 8 hours, we removed the male from the vial and presented females within a new male the following morning. After mating we removed males and transferred females to a new food vial and allowed females to lay eggs overnight. We removed females the following morning and kept oviposition vials at 25°C and counted the numbers of wild-type (focal) and brown (competitor) eyed offspring emerging 11 and 15 days later.

### Statistical analysis

Full details of the statistical analyses can be found in the online code repository (https://github.com/MartinGarlovsky/mito_age_fert). We performed statistical analysis in *R* v.4.2.2 (R Core Team 2022). We fit (generalised) linear mixed effects models ([G]LMMs) using *lme4* (Bates et al. 2015) and *lmerTest* (Kuznetsova et al. 2020). For fit LMMs with restricted estimates of maximum likelihood and used type-III ANOVA and Kenward-Roger’s F-tests and approximate denominator degrees of freedom. We checked model diagnostics using the *performance* (Lüdecke et al. 2021) and *DHARMa* (Hartig and Lohse 2022) packages. We checked model outputs to ensure the estimated degrees of freedom correctly reflected our experimental design, i.e., the number of replicate populations (N = 27). Estimated degrees of freedom were not always integer values due to how degrees of freedom are approximated for mixed models using the Kenward-Roger’s method (Halekoh and Højsgaard 2014). For GLMMs no denominator degrees of freedom are reported so we constructed comparable LMMs to check the appropriate degrees of freedom for the model. For all other models we performed type-III ANOVA tests using the ‘Anova’ function from the *car* package (Fox et al. 2023). We set options for contrasts to sum-to-zero and orthogonal polynomials. When interaction terms were not statistically significant, they were retained in models but not interpreted. We computed estimated marginal means (EMMs) and performed post-hoc tests corrected for multiple testing using the *emmeans* package (Lenth et al. 2023).

For the “geography based” analysis mitochondrial and nuclear genotype were defined by collection location of the founder populations, i.e., “A”, “B” or “C”. We included mitochondrial genotype, nuclear genotype, and the mitochondrial × nuclear interaction as fixed effects in all models and all two-way and three-way interactions for our fixed effects of interest (e.g., *age*, *sex*, etc.). Models followed the general form (in *R* notation):

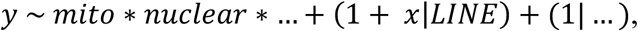

where y denotes the response variable of interest and ellipses denote other fixed (e.g., *age*, *sex*) or random (e.g., *block*, *individual identity*) effects included where applicable (detailed below). All models included mitonuclear population as a random intercept term and a random slope term (denoted by 1+ x in equation (1)) to allow each population to vary in its response.

For the analysis of body size, we scaled wing area to have mean zero and unit standard deviation and fit a LMM with sex as a random slope within population and family identity as a random intercept (see above).

Females showed age-specific reproductive periods and we therefore modelled these episodes separately (Novoseltsev et al. 2003). We modelled female early-life fecundity (summed progeny across the first 7 days: vials 1 and 2) using LMMs fit with the ‘lme’ function as visual inspection showed improved model diagnostic plots (as opposed to a GLMM with Poisson errors after accounting for overdispersion). For the analysis of female offspring production rate, we modelled the decline in offspring production for the period of decline (from vial 2 to vial 6) using a GLMM with Poisson errors and a log link, including the second order polynomial term for vial, and a random slope for vial within population and an observation level random effect (Harrison 2014).

We modelled male partner fecundity and sperm metabolic rate with LMMs with male identity as a random effect. Hatching success data showed distinct patterns in young vs. old males, and we therefore analysed young and old males separately. For young males we modelled hatching success with the response variable represented by a binomial vector consisting of the numbers of hatched (successes) and unhatched (failures) eggs using a GLMM with binomial errors and a logit link. In old males we modelled fertilisation success as a binary response indicating whether males sired any progeny, i.e., were fertile (1) or sterile (0) using a GLMM with binomial errors and a logit link. For analysis of sperm competition (P1 and P2), the response variable was represented by a binomial vector consisting of the numbers of wild-type (“successes”) and brown eyed (“failures”) offspring using GLMMs with binomial errors and a logit link. We included mating day and copulation duration as covariates, block and male identity as random effects, and an observation level random effect in binomial GLMMs to account for overdispersion (Harrison 2015). We modelled sperm viability in the control (t0) and stress treatment (t30) with the response variable represented by a binomial vector consisting of the numbers of live (“successes”) and dead (“failures”) sperm using GLMMs with binomial errors and a logit link with the numbers of live (“successes”) and dead (“failures”). We included male identity as a random effect and an observation level random effect. Sperm competition and sperm viability models returned a singularity warning message due to a zero-variance component estimate for the line random effect intercept. We retained these models after trying a range of alternatives and checking simplified models without the random slopes term that all resulted in the same qualitative findings for the fixed effects part of the model.

We modelled sperm quality with a LMM, using the mean difference in the proportion of viable sperm between t0 and t30 for each male as the response (Eckel et al. 2017). For the measurement of sperm viability, we collected all males on a single day for each block. However, the dissection, staining and incubation protocol necessitated measurements were carried out over several days. The measurement of sperm viability in young males (collected at 5 days old) thus extended for 8 days and that of old males (collected at 6 weeks old) for 6 days. To account for this variation in age we included dissection day (an extension of male age) and block as fixed covariates.

The initial experimental design based on geography allowed for a full factorial design. Sequencing of the mitochondrial genomes for all 27 mitonuclear population ((Vaught et al. 2020), Genbank NO. PRJNA532313) revealed 44 SNPs that differentiated between mitonuclear population by at least one site, resulting in 9 distinct mitochondrial haplotypes: mito-types 1 to 9 (Fig. 1). Therefore, to test differences associated with specific mitochondrial haplotypes we reanalysed the data replacing the mitochondrial genotype fixed effect (i.e., “A”, “B”, or “C”) with “mito-type” (mito-type 1 to 9). This led to the fixed effect model matrix being rank deficient (not every combination of mito-type × nuclear genotype was possible) with corresponding *R* warning messages. We inspected model diagnostic plots and outputs when fitting these models to ensure reasonable model estimates despite the unbalanced design.

In the results we present reaction norms of the model estimated marginal means for each mitochondrial genotype within each nuclear genotype. Plots of the raw data, including final sample sizes after excluding flies that died, escaped, produced no offspring (suggesting mating did not take place), or where females did not remate (for the sperm competition experiments) can be found in supplementary figures (Fig. S3-S11).

## RESULTS

### Body size

Body size showed a statistically significant effect of sex (LMM, F_1,18_ = 403.05, *p* < 0.001) and nuclear genotype (F_2,18_ = 10.57, *p* < 0.001) but no interaction (Fig. 2). Females were larger than males in all comparisons (post-hoc Tukey’s HSD; all *p* < 0.001) as expected in *Drosophila*. Flies with the A nuclear genotype were smaller than B (post-hoc Tukey’s HSD; *p* = 0.003) and C flies (*p* = 0.002) but B and C did not differ (*p* = 0.988).

**Figure 2.**
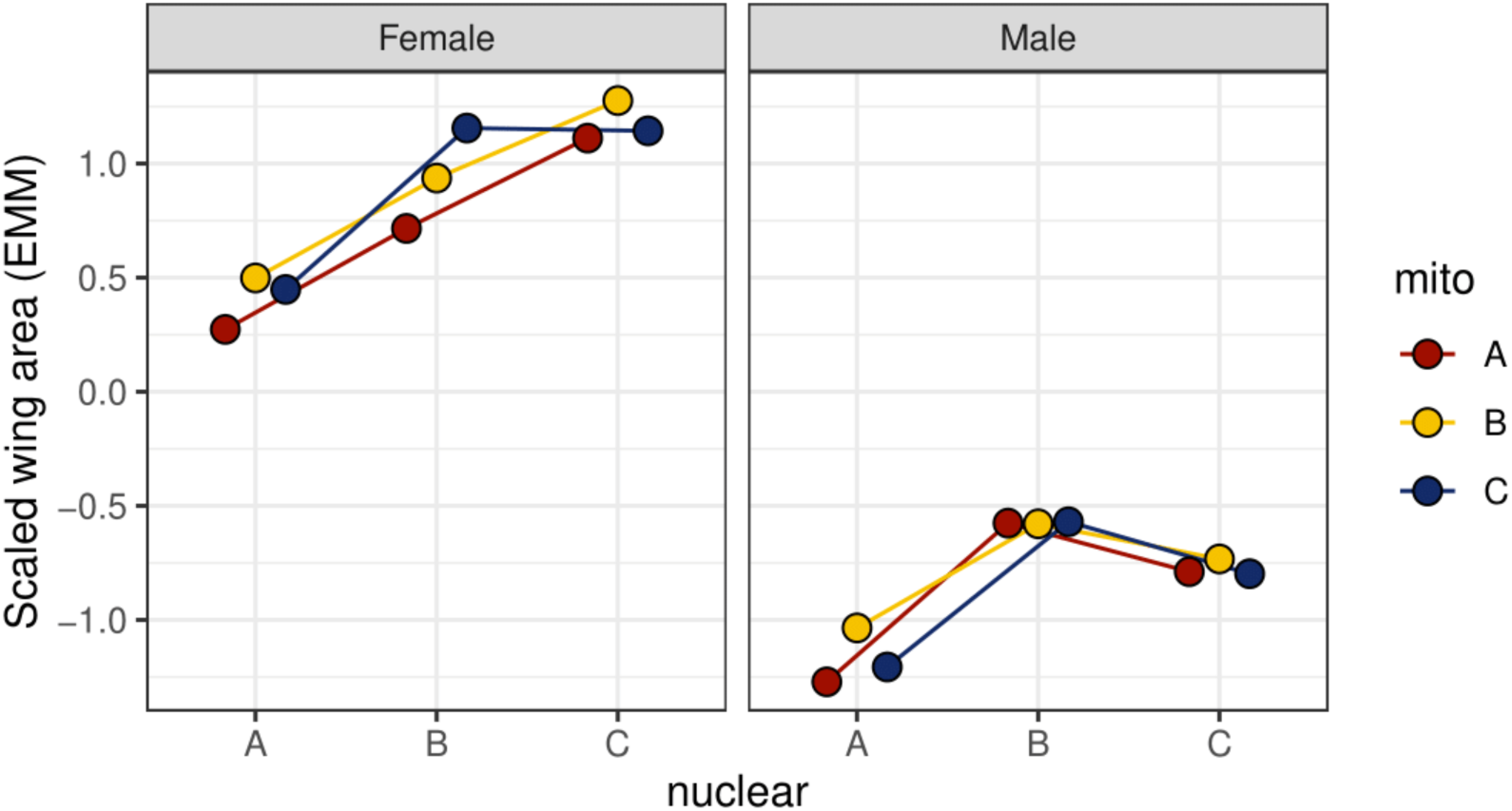
Body size (scaled wing area) differences between experimentally matched and mismatched mitonuclear genotypes in females (left) and males (right) *Drosophila melanogaster*. Points show estimated marginal means (EMM) for each mitochondrial genotype (“mito”) within each nuclear genotype (x-axis).

### Female reproductive success

#### Early-life female fecundity

There was a statistically significant effect of nuclear genotype on early-life female fecundity (LMM, F_2,17.77_ = 18.27, *p* < 0.001; Fig. 3a). Females with the A nuclear genotype produced more offspring during the first seven days after mating (first two vials) compared to B or C females (post-hoc Tukey’s HSD, A vs. B: *p* < 0.001; A vs. C: *p* < 0.001), whereas B and C females did not differ (*p* = 0.728).

**Figure 3.**
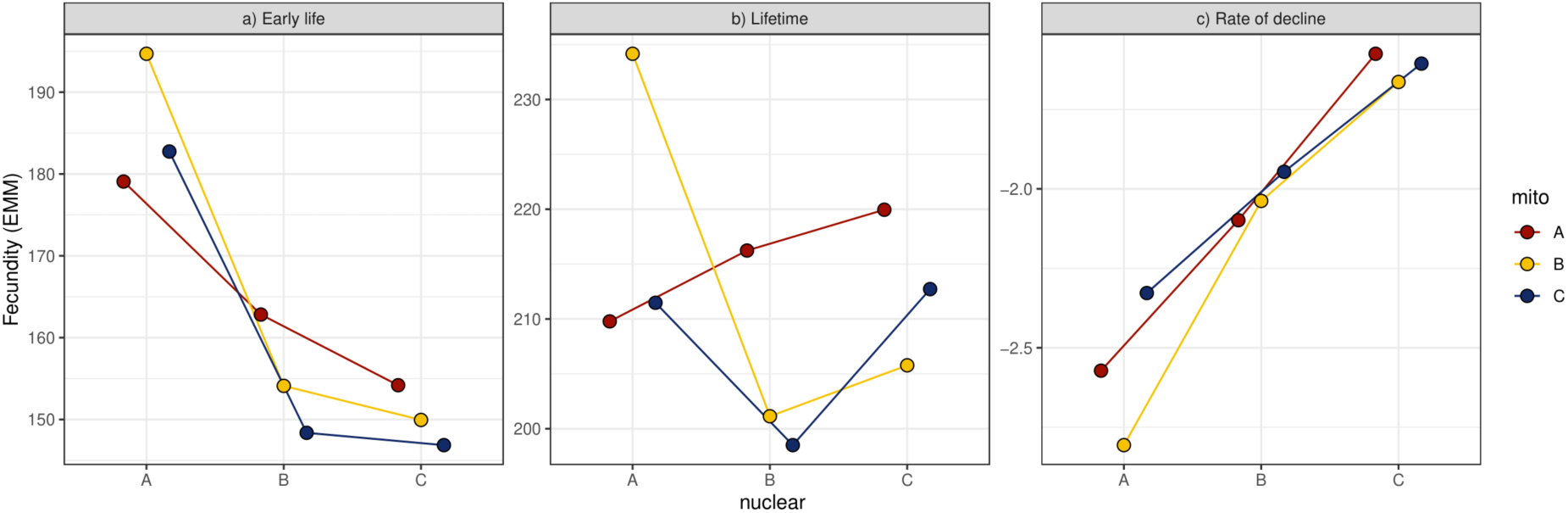
Female fitness parameters of experimentally matched and mismatched mitonuclear genotypes in female *Drosophila melanogaster*. a) Early life fecundity (first 7 days, numbers of offspring produced pooled across first two vials). b) Lifetime fecundity (numbers of progeny produced over 21 days). c) Rate of offspring production (negative values indicate a faster rate of decline) from a model of female per-vial offspring production from the onset of reproductive senescence (excluding first and last vials: 1 and 7; see *Methods*). Points show estimated marginal means (EMM) for each mitochondrial genotype (“mito”) within each nuclear genotype (x-axis). Note different y-axis scales in a), b), and c).

#### Lifetime female fecundity

There was no effect of mitochondrial genotype, nuclear genotype, or the mitonuclear interaction on female lifetime fecundity (Fig. 3b), in line with previous results from the same mitonuclear populations measured under benign conditions (Dobson et al. 2023). Females produced an average of 212 offspring (95% confidence intervals [CIs]: 205 – 219) over the course of the experiment (21 days) after a single mating.

#### Decline in offspring production

The numbers of offspring produced declined over female lifespan (between vials 2 and 6; see *Methods*; Fig. S4) and the rate of decline differed between females based on nuclear genotype (nuclear genotype × vial interaction, Poisson GLMM, X^2^ = 72.65, df = 2, *p* < 0.001). The rate of decline differed between all nuclear genotypes (post-hoc Tukey’s HSD, all *p* < 0.001). Females with the A nuclear genotype had the fastest rate of decline, likely resulting from a higher early life fecundity, while females with the C nuclear genotype had the slowest rate of decline (Fig. 3c). There was no effect of mitochondrial genotype or the mitonuclear interaction.

We also modelled female reproductive senescence using survival analysis, based on the onset of infertility, i.e., the time (in vials) at which females stopped producing any progeny which produced the same qualitative results (see online supplementary material).

### Male reproductive success

#### Sperm viability

The initial proportion of live sperm (t0) showed a statistically significant nuclear genotype × age interaction (binomial GLMM, X^2^ = 13.10, df = 2, *p* = 0.001; Fig. 4a - circles). Old males with the A or C nuclear genotype had lower sperm viability than young males (post-hoc Tukey’s HSD, A nuclear young vs. old, *p* < 0.001, C nuclear young vs. old, *p* = 0.005), whereas B males did not show an effect of age (*p* = 0.163). In young males there was no difference in sperm viability between nuclear genotypes (all *p* > 0.257). However, in old age, A males had lower sperm viability than B males (*p* = 0.001) and C males (*p* < 0.001), but B and C males did not differ (*p* = 0.984).

**Figure 4.**
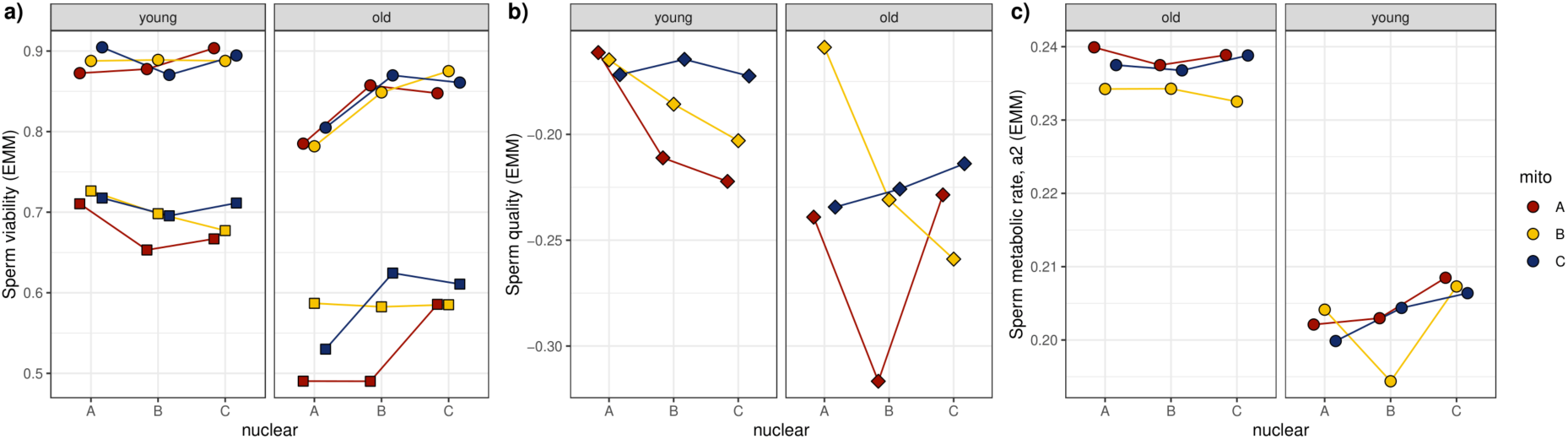
Sperm performance parameters in young (5 day old) and old (6-week-old) males. a) Sperm viability at time t0 (circles) and after 30 min in stressor medium (t30; squares) averaged over blocks and days. b) Sperm quality, i.e., the difference in sperm viability between t0 and t30. c) Sperm metabolic rate (the proportion of protein bound NAD(P)H, a2). Points show estimated marginal means (EMM) for each mitochondrial genotype (“mito”) within each nuclear genotype (x-axis). Note different y-axis scales.

At t30, the proportion of live sperm showed a statistically significant effect of mitochondrial genotype (binomial GLMM, X^2^= 7.12, df = 2, *p* = 0.028; Fig. 4a - squares), age (X^2^= 58.10, df = 1, *p* < 0.001), block (X^2^= 15.12, df = 1, *p* < 0.001) and the nuclear genotype × age interaction (X^2^= 6.02, df = 1, *p* = 0.049). Males with the A mitochondrial genotype had lower sperm viability than C males (post-hoc Tukey’s HSD, *p* = 0.042), old males had lower viability than young males (*p* < 0.001), and sperm viability was higher in the first block compared to the second (*p* < 0.001).

#### Sperm quality

On average, sperm viability declined by 20% after the 30-minute incubation period in stressor medium. Sperm quality, i.e. the temporal decline in sperm viability between t0 and t30 (Eckel et al. 2017) showed a statistically significant effect of age (LMM, F_1,22.64_ = 9.31, *p* = 0.006), block (F_1,579.2_ = 10.20, *p* = 0.001), and dissection day (F_1,576.5_ = 7.55, *p* = 0.006; Fig. 4b). Old males had lower sperm quality than young males (post-hoc Tukey’s HSD, *p* = 0.006), sperm quality was higher in the first block compared to the second (*p* = 0.002), and sperm quality declined with dissection day (β = −0.010, 95% CIs: −0.018, −0.003).

#### Sperm metabolic rate

Sperm metabolic rate showed a statistically significant effect of male age (LMM, F_1,17.99_ = 240.34, *p* < 0.001; Fig. 4c) but no effect of mitochondrial or nuclear genotype. Old males’ sperm had higher metabolic rates (post-hoc Tukey’s HSD; *p* < 0.001). This result was robust to inclusion or exclusion of an extreme outlier (Fig. S3) and confirmed previous results of increased sperm metabolic rates in older male *Drosophila melanogaster* (Turnell & Reinhardt 2020).

#### Partner fecundity

The numbers of eggs laid by tester females after mating with a mitonuclear population male showed a statistically significant three-way mito × nuclear × age interaction (LMM, F_4,36.09_ = 3.51, *p* = 0.016; Fig. 5a). Young males had higher fecundity than old males in all comparisons (post-hoc Tukey’s HSD, all *p* < 0.001). In younger males, mismatched BC males had higher fecundity than coevolved CC males (*p* = 0.023) and BA males (*p* = 0.006). In old age, coevolved BB males had lower fecundity than mismatched BA males (*p* = 0.010).

**Figure 5.**
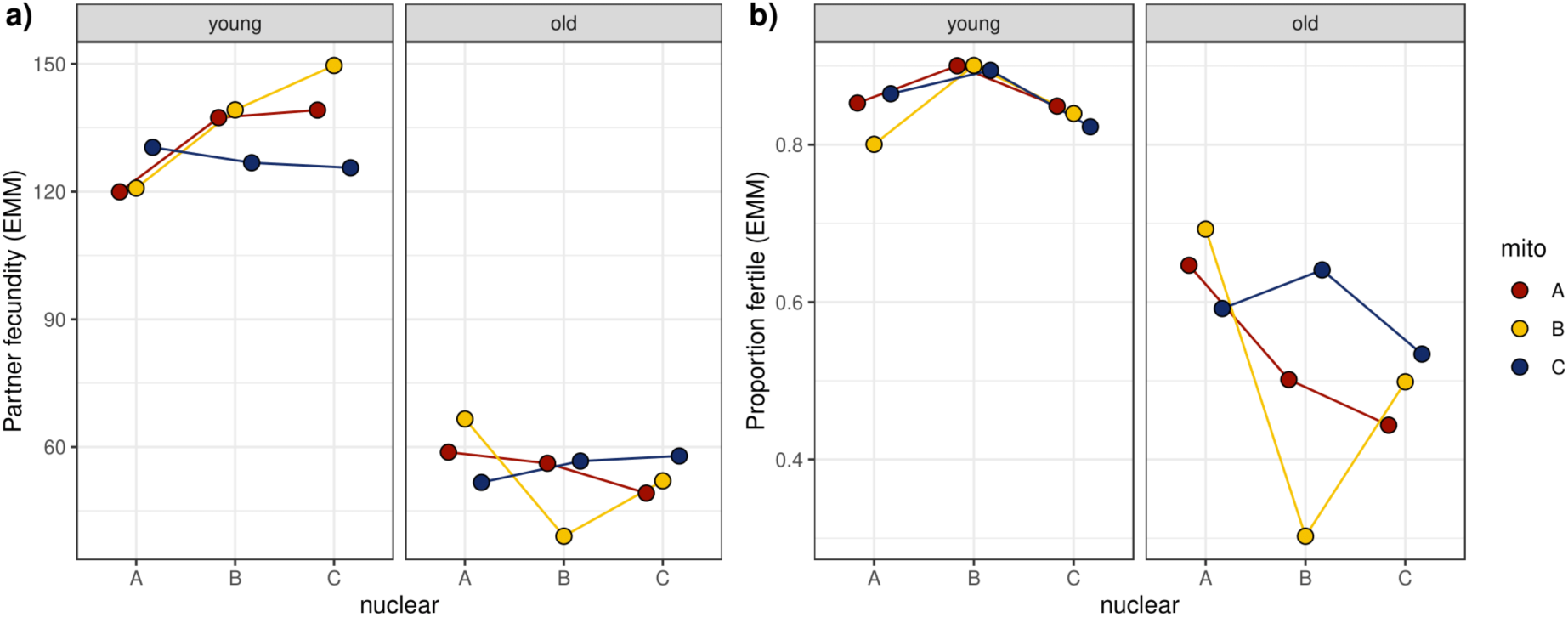
Male fitness parameters in young (5 days old) and old (6-week-old) males. a) Male partner fecundity (numbers of eggs laid by standard tester female) for young (left) and old (right) males. b) Fertility of young males (left, the proportion of eggs laid by tester females that hatched) and old males (right, probability of sterility). Points show estimated marginal means (EMM) for each mitochondrial genotype (“mito”) within each nuclear genotype (x-axis). Note different y-axis scales.

#### Male fertilisation success

In young males, there was a statistically significant effect of nuclear genotype on the proportion of hatching eggs laid by tester females (binomial GLMM, X^2^ = 26.19, df = 2, *p* < 0.001; Fig. 5b). Males with the B nuclear genotype had higher hatching success than A males (post-hoc Tukey’s HSD, *p* < 0.001) and C males (*p* < 0.001), whereas A and C males did not differ (*p* = 0.969).

In old males, we modelled hatching success as a binary response indicating whether males were fertile (any eggs hatched) or sterile (no eggs hatched). There was a statistically significant effect of nuclear genotype on the probability of sterility in old males (binomial GLMM; X^2^ = 6.82, df = 2, *p* = 0.033; Fig. 5c). B males had a marginally higher probability of sterility than A males (post-hoc Tukey’s HSD; *p* = 0.059).

#### Sperm competition

Sperm defence (P1) showed a statistically significant effect of nuclear genotype (binomial GLMM, X^2^ = 12.79, df = 2, *p* = 0.002) and mating day (X^2^= 16.86, df = 1, *p* < 0.001). Males with nuclear genotype B fared worse in sperm defence than males with the A (post-hoc Tukey’s HSD, *p* = 0.001) or C (*p* = 0.036) nuclear genotype, whereas A and C males did not differ (*p* = 0.503) (Fig. 6). Males whose partner remated on the second day had lower P1 than when remating took place on the first day (*p* < 0.001).

**Figure 6.**
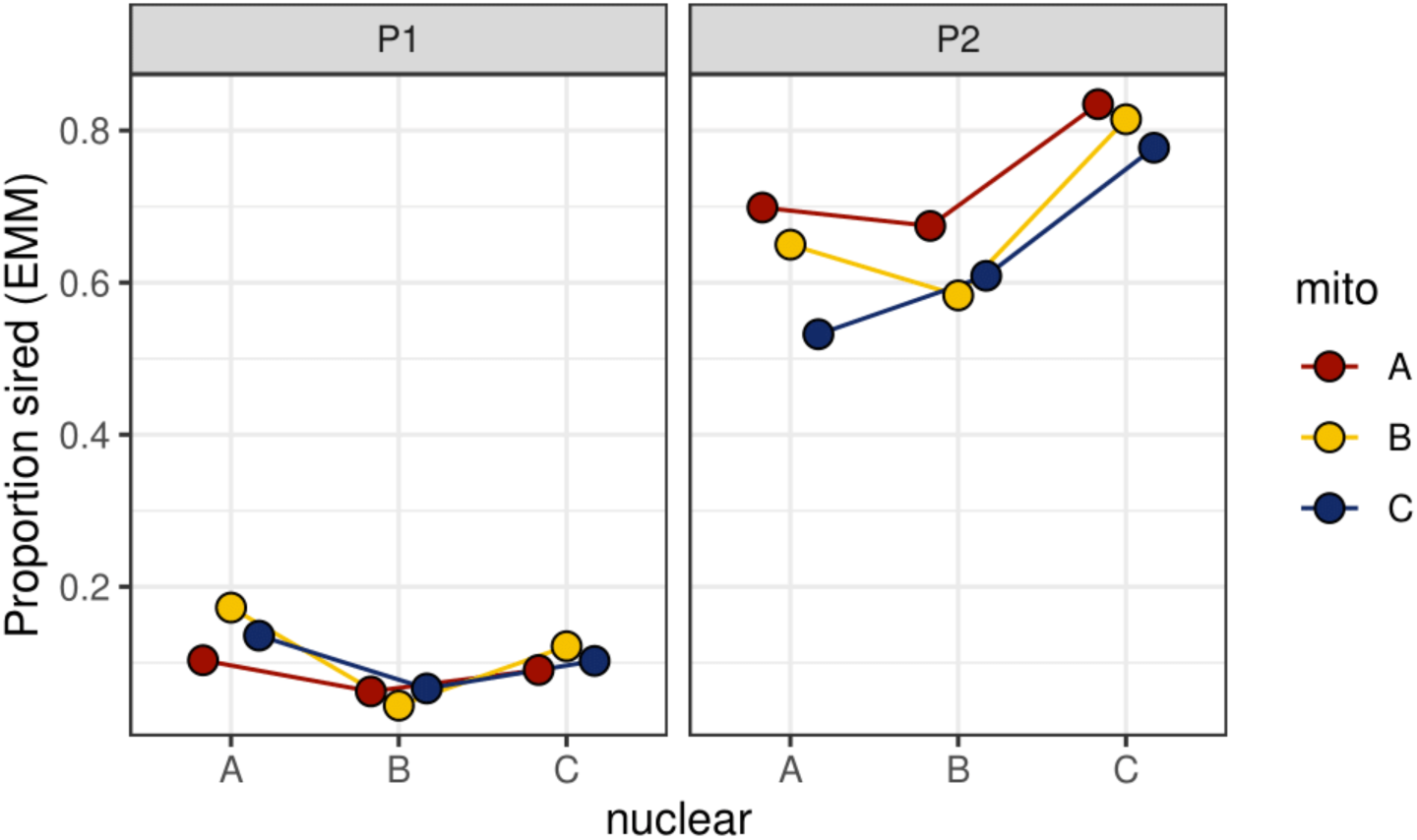
Sperm competition success in young males (aged 5-6 days). Sperm defence, (P1, left) and sperm offence (P2, right) for males competing against isogenic tester (*bw*) males. Points show estimated marginal means (EMM) for each mitochondrial genotype (“mito”) within each nuclear genotype (x-axis) averaged across mating days.

Sperm offence (P2) showed a statistically significant effect of mitochondrial genotype (binomial GLMM, X^2^ = 10.68, df = 2, *p* = 0.005), nuclear genotype (X^2^= 54.52, df = 2, *p* < 0.001), and mating day (X^2^= 49.92, df = 1, *p* < 0.001) but no effect of the mito × nuclear interaction (Fig. 6). Males with mitochondrial genotype A had higher P2 than C males (post-hoc Tukey’s HSD, *p* = 0.003), while A and B (*p* = 0.180) and B and C (*p* = 0.311) males did not differ. Males with C nuclear genotype had higher P2 than A (*p* < 0.001) or B (*p* < 0.001) males, but A and B males did not differ (*p* = 0.980). Males whose partner remated on the second day had lower P2 compared to those remating on the first day (*p* < 0.001).

There was no correlation between P1 and P2 (model estimated marginal means for each mitonuclear genotype; Spearman’s rank correlation, ρ = 0.05, S = 114, *p* = 0.91), indicating the mitonuclear genotype with best/worst defensive paternity success was not the best/worst in an offensive role (Fig. S12).

### Mito-type analysis

Finally, we reanalysed the data using the 9 major mitochondrial haplotypes – “mito-types” (Fig. 1). None of the female traits (early life and lifetime fecundity, and reproductive senescence) or male traits (sperm viability, sperm quality, sperm metabolic rate, partner fecundity or hatching success, and sperm competitive success) showed an effect of mito-type or the mito-type × nuclear genotype interaction.

#### Body size

Body size showed a statistically significant mito-type × nuclear genotype × sex interaction (LMM: F_7,9_ = 6.73, *p* = 0.005) indicating sex-specific mitonuclear epistasis (Fig. S13). In a B nuclear background, mito-type 1 (A-like) females were smaller than females with mito-type 3 and mito-type 4 (A-like), mito-type 7 (present in both B and C populations) and mito-type 8 (C-like) (post-hoc Tukey’s HSD; all *p* < 0.041). Mito-type 1 and mito-type 3 differ at a single SNP at position 12512 (G/A) in NADH-ubiquinone oxidoreductase chain 1 (*ND1*; FBgn0013679), which results in a non-synonymous substitution of proline to serine at codon 50, whereas mito-type 4 differs from mito-type 1 in 6 SNPs but not position 12512 (Fig. 1).

Likewise, mito-types 7 and 8 differ from mito-type 1 at many SNPs but crucially both haplotypes have a G at position 12512 (as do all other haplotypes and the FlyBase.org reference allele). Body size of mito-type 1 females was smaller in B compared to C nuclear backgrounds (*p* = 0.005), indicating a specific interaction of this “A-like” mitochondrial SNP at position 12512 in a B nuclear background. Together, this implicates the G/A polymorphism at position 12512 as a negative epistatic interaction between A mitochondrial and B nuclear effecting female body size. Females with mito-type 7 (present in both B and C populations) were also smaller in A nuclear compared to coevolved C nuclear background (*p* = 0.048), again suggesting an incompatibility of the B/C-like mitochondria in the A nuclear background.

In males we also found a mito-type × nuclear genotype interaction effect on body size involving mito-types 1, 3, 7 and 8, again implicating the interaction between the SNP at position 12512 in *ND1* in different nuclear backgrounds. In an A nuclear background males were smaller than B nuclear males with mito-type 7 (*p* = 0.005) and mito-type 8 (*p* = 0.041). Likewise, males with an A nuclear background were smaller than C nuclear males with mito-type 1 (*p* = 0.032) and mito-type 7 (*p* = 0.046). Flies with mito-type 1 showed no sex difference in body size in the B nuclear background (*p* = 0.945).

#### Partner fecundity

The numbers of eggs laid by tester females mated to mitonuclear population males showed a statistically significant three-way mito-type × nuclear × age interaction (LMM, F_1,18.04_ = 6.40, *p* = 0.002). In young males, mito-type 7 (B/C-like) sired fewer offspring in a novel A compared to co-evolved C nuclear background (post-hoc Tukey’s HSD, *p* = 0.043) (Fig. S14). In old males, mito-type 6 had higher fecundity in a novel A compared to a co-evolved B nuclear background (*p* = 0.046). Mito-type 6 differs from the other mitochondrial haplotype found in these comparisons (mito-type 7) at a single C/T polymorphism at position 13934 in *mt:lrRNA* encoding the 16S rRNA of the mitochondrial ribosome, previously shown to be involved in diet × mitonuclear effects on fitness (Dobson et al. 2023).

### Coevolved analysis

Finally, we tested whether there was an effect of matched (“coevolved”) vs. mismatched mitonuclear combinations but found no overall effect of matched vs. mismatched genotypes or the interaction with sex or age for any parameter we measured.

## DISCUSSION

Mitonuclear mismatch is predicted to have significant fitness consequences by disturbing cooperation between coevolved mitochondrial and nuclear encoded components of the electron transport system (Dobler et al. 2018). Negative effects of mismatched mitonuclear genotypes are expected to be most prominent in males due to the sex-biased mutation load arising from the maternal inheritance of mitochondria (Dowling and Adrian 2019). Here we tested the effects of mitochondrial replacement on female and male fitness and how these effects change with age. We found no overall difference in fitness for matched compared to mismatched mitonuclear combinations in females or males. We also found no effect of mitonuclear epistasis on female fitness parameters. Assuming that our matched combinations are coevolved, the lack of mitonuclear epistasis effect in females suggests that females were relatively robust to perturbations of mitonuclear coevolution. We did find an effect of mitonuclear epistasis in males, affecting the fecundity of their mating partners. However, the effect was not in the direction predicted by cooperative coevolution between interacting genomes.

### Mitonuclear epistasis

Mitochondrial replacement revealed an age-dependent effect of mitonuclear epistasis in just three out of 36 possible comparisons. In young males, mismatched BC males had higher reproductive success (partner fecundity) than coevolved CC males and BA males. In old age, mismatched BA males had higher reproductive success than coevolved BB males. Idiosyncratic mitonuclear effects have frequently been reported in male traits (Reinhardt et al. 2013; Dobler et al. 2014, 2018; Immonen et al. 2016a). Sex-specific fitness effects of mitochondrial replacement might be expected in males if novel mitonuclear interactions “release” nuclear alleles from a male deleterious mitochondrial allele that has fixed within populations due to the Muller’s ratchet. Alternatively, the B mitochondrial haplotype may not harbour male deleterious alleles, but our experimental mitonuclear pairing resulted in a male specific effect on late life reproduction. Currently it is not clear whether the possible beneficial effect occurred because the laboratory B population was inbred or whether inbreeding is already found in wild B populations. Should beneficial mitonuclear mismatch be common in nature, it might represent an important counterforce to the mother’s curse (Unckless and Herren 2009; Wade and Brandvain 2009; Hedrick 2012).

Dissecting the mitochondrial genotype further based on the major allele frequencies within each mitonuclear population revealed specific mitochondrial SNPs or haplotypes involved in mitonuclear mismatch with both positive and negative fitness effects. The mito-type analysis revealed mitonuclear epistasis effecting female and male body size. The G/A polymorphism at position 12512 in *ND1* affecting body size encodes a non-synonymous mutation (proline to serine) (Fig. 1). Mitochondrial variation in complex 1 of the electron transport system (which includes *ND1*) was recently shown to result in a sex-specific incompatibility in a mismatched nuclear background; causing reduced female survival, fertility, and respiratory rate, increased H_2_O_2_ flux and loss of redox control (Camus et al. 2023).

For male partner fecundity, the mito-type analysis implicated mito-type 6 in old males as the causal haplotype that is unfavourable in a coevolved (Beninese) nuclear background. The single SNP differentiating mito-type 6 from the other mito-type found in Beninese nuclear backgrounds (mito-type 7) is a C/T polymorphism at position 13934 in *mt:lrRNA* encoding the 16S rRNA of the mitochondrial ribosome (Fig. 1). We found the Thymine allele was associated with poorer fecundity in a Beninese nuclear background in males. Although the effects of this SNP on components of reproduction were male- and age-specific in this study, previous evidence suggests that there are conditions under which this same SNP manifests effects on both sexes (Dobson et al. 2023). Mitonuclear interactions involving this same C/T polymorphism were recently shown to affect female progeny production, fertilisation success and development in response to diet (Dobson et al. 2023). Females carrying the Thymine allele showed reduced fertilisation success in both the Australian and Beninese nuclear background on an amino acid-enriched diet, whereas females with the Cytosine allele had lower fertility only in the Australian nuclear background (Dobson et al. 2023). The Cytosine allele impaired fertility after feeding on an amino acid-enriched diet and on lipid-only diet in both Australian and Beninese nuclear background but the Thymine allele did so only in the Australian and not the Beninese background (Dobson et al. 2023). Thus, the effects associated with this SNP do not seem to align with expectations under the mother’s curse hypothesis.

Several previous studies found mitonuclear incompatibility affecting female fitness (e.g., Dowling et al. 2008, Dordević et al. 2017; Immonen et al. 2016a; Zhang et al. 2017, Vaught et al. 2020, Camus et al. 2023). By contrast, we found no overall effect of mitonuclear interactions on female fitness. This result is similar to previous evidence from the same mitonuclear populations measured under similar conditions (Dobson et al. 2023), as well as to other studies reporting little evidence for mitonuclear mismatch causing variation in female fitness (Keaney et al. 2020). These studies suggest that females may be relatively insensitive to effects of novel mitonuclear interactions or that nucleotide divergence between haplotypes in *D. melanogaster* is too small to generate negative epistatic effects. Another non-mutually exclusive possible explanation for the differences in results between previous studies and those reported here is due to differences in experimental design to create the mitochondrial replacement panels. Our experimental design aimed to preserve segregating mitochondrial and nuclear to reflect variation in populations found in nature, whereas most other studies use isogenic strains with fixed mitochondrial and nuclear variants.

### Independent mitochondrial and nuclear effects

In males we found mitochondrial genotype alone caused variation in male-specific traits; sperm quality and sperm offensive paternity success (P2). Mitochondrial variation has frequently been invoked to explain variation in male traits (Smith et al. 2010; Patel et al. 2016; Martikainen et al. 2017; Vaught and Dowling 2018), sometimes showing mitonuclear variation e.g., (Clancy et al. 2011; Innocenti et al. 2011; Yee et al. 2013), unlike in our study. We also found nuclear genotype independently caused variation in body size in both sexes (A nuclear genotype flies were smaller than B or C flies), and female traits (early-life fecundity and reproductive rate) and male traits (sperm viability, male fertilisation success, and sperm competition success). Females with the A nuclear genotype produced more offspring in the first week after mating but also experienced earlier reproductive senescence. Conversely, we found no differences between nuclear genotypes in female lifetime reproductive success. This suggests a different solution between nuclear genotypes posed by the trade-off between early vs. later reproductive output, ultimately resulting in similar lifetime reproductive success for all three populations surveyed. In nature these differences in investment may lead to differences in fitness, for instance, in environments where early mortality would favour early investment (Stearns 1992).

### Ageing effects

We found 6-week-old males performed more poorly than young males (5 days old) in every parameter we measured (sperm viability and quality, partner fecundity and fertilisation success). Previous studies have shown that sperm traits and fitness decrease with male age (Cornwallis et al. 2014; Immonen et al. 2016a; Fricke and Koppik 2019; Sepil et al. 2020) but a recent meta-analysis (including the data presented here) reject this observation as a general pattern (Sanghvi et al. 2024). The reduced performance of sperm parameters in old age was accompanied by higher sperm metabolic rates. In *D. melanogaster*, sperm from older males with higher sperm metabolic rate had accumulated more ROS but sperm produced them at a lower rate (Turnell & Reinhardt 2020). Genetic engineering to suppress ROS production in the germline showed increased partner fecundity and a longer fertile period (Turnell et al. 2021). Our data suggest that mitochondrial genotype variation seemingly did not affect sperm metabolism and that any previously identified strategies to reduce sperm ROS by older males (Turnell & Reinhardt 2020) are insufficient to outweigh the deleterious effects of ageing.

In conclusion, we found evidence for mitonuclear epistasis affecting several traits in both females and males, but no overall effect of mitonuclear mismatch. The direction of effects was often not that expected if mismatched mitonuclear interactions are deleterious. Novel combinations of mitochondrial and nuclear genotypes more often resulted in beneficial outcomes, suggesting heterosis and the potential escape from Muller’s ratchet. We were able to identify two candidate SNPs in the mitochondrial genome associated with mitonuclear fitness interactions, one of which has previously been shown to affect female fitness. We also found ageing had a negative effect on several male traits linked to fitness and showed this was linked to faster metabolic rates in the sperm of old males. This study provides a detailed analysis of the fitness consequences of mitonuclear interactions in males. Our results may more generally help to explain idiosyncratic and often unpredictable outcomes of mitonuclear interactions on male fitness. Overall, the results suggest that males are more sensitive to both main effects of mitochondrial variation (affecting sperm traits), and to effects of mitonuclear interactions, relative to females. Given the maternal inheritance of the mitochondrial genome and predicted female-specific selection on the genome, this may suggest that this mitochondrial variation seen in males is associated with a sex specific mutation load (mother’s curse).

## DATA AVAILABILITY

Data will be archived at Dryad: 10.5061/dryad.dv41ns27n. All code will be deposited on *GitHub*: https://martingarlovsky.github.io/mito_age_fert/.

## AUTHOR CONTRIBUTIONS

RD and KR conceived the study; MDG, RG, RD, and KR wrote the manuscript; DKD and RD designed and created the strains; RG carried out the experiments, supervised by KR, RD; MDG, RG, SV, and RD analysed the data. MDG and KR wrote the first version of the manuscript, and all authors contributed feedback and editing of subsequent drafts. All authors approved the final version of the manuscript.

## ACKNOWLEDGEMENTS

We thank Christin Froschauer, Susanne Broschk, Cornelia Thodte, Wei Dong and Emmely Voigt for help in the lab and maintaining the introgression lines and John Jackson and Adam Dobson for helpful discussion about statistical analysis. Ensieh Habibi carried out the sperm competition trials. Jonci Wolff provided the mitochondria enrichment protocol; the Biotec TU Dresden (Deep Sequencing Group - SFB655) sequenced the mitochondrial genome. Cornelia Wetzker provided FLIM training to RG.

